# The great tit HapMap project: a continental-scale analysis of genomic variation in a songbird

**DOI:** 10.1101/561399

**Authors:** Lewis G. Spurgin, Mirte Bosse, Frank Adriaensen, Tamer Albayrak, Christos Barboutis, Eduardo Belda, Andrey Bushuev, Jacopo G. Cecere, Anne Charmantier, Mariusz Cichon, Niels J. Dingemanse, Blandine Doligez, Tapio Eeva, Kjell Einar Erikstad, Vyacheslav Fedorov, Matteo Griggio, Dieter Heylen, Sabine Hille, Camilla A. Hinde, Elena Ivankina, Bart Kempenaers, Anvar Kerimov, Milos Krist, Laura Kvist, Veronika N. Laine, Raivo Mänd, Erik Matthysen, Ruedi Nager, Boris P. Nikolov, A. Claudia Norte, Markku Orell, Jenny Ouyang, Gergana Petrova-Dinkova, Heinz Richner, Diego Rubolini, Tore Slagsvold, Vallo Tilgar, János Török, Barbara Tschirren, Csongor I. Vágási, Teru Yuta, Martien A.M. Groenen, Marcel E. Visser, Kees van Oers, Ben C. Sheldon, Jon Slate

**Affiliations:** School of Biological Sciences, University of East Anglia, Norwich Research Park, United Kingdom; Edward Grey Institute, University of Oxford, United Kingdom; Department of Animal Ecology, Netherlands Institute of Ecology (NIOO-KNAW), Wageningen, the Netherlands; Animal Breeding and Genomics Centre, Wageningen University, the Netherlands; Department of Ecological Science, Animal Ecology Group, Vrije Universiteit Amsterdam, Amsterdam, The Netherlands; Evolutionary Ecology Group, Department of Biology, University of Antwerp, Universiteitsplein 1, 2610 Antwerp, Belgium; Department of Biology, Mehmet Akif Ersoy University, Science and Art Faculty, Ortulu, Burdur, Turkey; Hellenic Ornithological Society / BirdLife Greece, Themistokleous 80, GR-10681 Athens; Institut d’Investigació per a la Gestió Integrada de Zones Costaneres, Campus de Gandia, Universitat Politècnica de València, Carrer Paranimf 1, E-46730 Grau de Gandia (València), Spain; Faculty of Biology, Lomonosov Moscow State University, Moscow 119234, Russia; Area Avifauna Migratrice, Istituto Superiore per la Protezione e la Ricerca Ambientale (ISPRA), via Ca’ Fornacetta 9, I-40064, Ozzano Emilia, (BO), Italy; CEFE-CNRS, UMR 5175, 1919, route de Mende, F34293 Montpellier Cedex 5, France; Institute of Environmental Sciences, Jagiellonian University, Gronostajowa 7, 30-387 Kraków, Poland; Behavioural Ecology, Department of Biology, Ludwig Maximilians University of Munich, Planegg-Martinsried, Germany; UMR CNRS 5558—LBBE, Biométrie et Biologie Évolutive, UCB Lyon 1 - Bât. Grégor Mendel, 43 bd du 11 novembre 1918, 69622 VILLEURBANNE cedex, France; Department of Ecology and Evolution, Animal Ecology, Evolutionary Biology Centre, Uppsala University, Sweden; Department of Biology, University of Turku, Turku 20014, Finland; Norwegian Institute for Nature Research, FRAM-High North Research Centre for Climate and the Environment, 9296 Tromsø, Norway; Department of Biology, University of Padova, Via U. Bassi 58/B, I-35131, Padova, Italy; Department of Ecology and Evolutionary Biology, Princeton University, Princeton, NJ, United States of America; Interuniversity Institute for Biostatistics and statistical Bioinformatics, Hasselt University, Diepenbeek, Belgium; Institute of Wildlife Biology and Game Management, University of Natural Resources and Life Science, A-1180 Vienna, Austria; Behavioural Ecology Group, Department of Life Sciences, Anglia Ruskin University, East Road Cambridge, Cambridgeshire, CB1 1PT, UK; Zvenigorod Biological Station of Lomonosov Moscow State University, P.O. Box Shikhovo, Odintsovo District, Moscow 143036, Russia; Max Planck Institute for Ornithology, Department of Behavioural Ecology & Evolutionary Genetics, Eberhard-Gwinner-Straße, House 5, 82319 Seewiesen (Starnberg), Germany; Department of Zoology and Laboratory of Ornithology, Faculty of Science, Palacký University, Olomouc 77147, Czech Republic; Department of Ecology and Genetics, P.O.Box 3000, 90014-University of Oulu, Finland; Finnish Museum of Natural History, University of Helsinki, FI-00014, Finland; Department of Zoology, Institute of Ecology and Earth Sciences, University of Tartu, Vanemuise 46, Tartu 51014, Estonia; Institute of Biodiversity, Animal Health and Comparative Medicine, University of Glasgow, Glasgow G12 8QQ, United Kingdom; Bulgarian Ornithological Centre, Institute of Biodiversity and Ecosystem Research, Bulgarian Academy of Sciences, 1 Tsar Osvoboditel Blvd, 1000 Sofia, Bulgaria; MARE - Marine and Environmental Sciences Centre, Department of Life Sciences, Faculty of Sciences and Technology, University of Coimbra, Portugal; University of Nevada, Reno, NV, United States of America; Evolutionary Ecology Lab, Institute of Ecology and Evolution, University of Bern, Bern 3012, Switzerland; Dipartimento di Scienze e Politiche Ambientali, Università degli Studi di Milano, via Celoria 26, I-20133, Milano, Italy; Centre for Ecological and Evolutionary Synthesis (CEES), Department of Biosciences, University of Oslo, P.O. Box 1066, Blindern, 0316 Oslo, Norway; Behavioural Ecology Group, Department of Systematic Zoology and Ecology, Eötvös Loránd University, Budapest H-1117, Hungary; University of Exeter, Centre for Ecology and Conservation, Treliever Road, Penryn, TR10 9FE, United Kingdom; Evolutionary Ecology Group, Hungarian Department of Biology and Ecology, Babeş-Bolyai University, Cluj-Napoca, Romania; Behavioural Ecology Research Group, Department of Evolutionary Zoology, University of Debrecen, Debrecen, Hungary; Graduate School of Environmental Science, Hokkaido University, N10 W5 Sapporo, Hokkaido 060-0810, Japan; Groningen Institute for Evolutionary Life Sciences (GELIFES), University of Groningen, Groningen, the Netherlands; Department of Animal and Plant Sciences, University of Sheffield, United Kingdom

## Abstract

A major aim of evolutionary biology is to understand why patterns of genomic diversity vary among populations and species. Large-scale genomic studies of widespread species are useful for studying how the environment and demographic history shape patterns of genomic divergence, and with the continually decreasing cost of sequencing and genotyping, such studies are now becoming feasible. Here, we carry out one of the most geographically comprehensive surveys of genomic variation in a wild vertebrate to date; the great tit (*Parus major*) HapMap project. We screened *ca* 500,000 SNP markers across 647 individuals from 29 populations, spanning almost the entire geographic range of the European great tit subspecies. We found that genome-wide variation was consistent with a recent colonisation across Europe from a single refugium in South-East Europe, with bottlenecks and reduced genetic diversity in island populations. Differentiation across the genome was highly heterogeneous, with clear “islands of differentiation” even among populations with very low levels of genome-wide differentiation. Low local recombination rate in the genome was a strong predictor of high local genomic differentiation (*F*_*ST*_), especially in island and peripheral mainland populations, suggesting that the interplay between genetic drift and recombination is a key driver of highly heterogeneous differentiation landscapes. We also detected genomic outlier regions that were confined to one or more peripheral great tit populations, most likely as a result of recent directional selection at the range edges of this species. Haplotype-based measures of selection were also related to recombination rate, albeit less strongly, and highlighted population-specific sweeps that likely resulted from positive selection. These regions under positive selection contained candidate genes associated with morphology, thermal adaptation and colouration, providing promising avenues for future investigation. Our study highlights how comprehensive screens of genomic variation in wild organisms can provide unique insights into evolution.

## Introduction

Since the first studies of allozyme variation in humans [1] and *Drosophila* [2,3], there has been great interest in explaining how evolutionary and ecological processes shape the patterns of genetic variation observed within and among natural populations. One focus of research and debate in this area has been on quantifying the roles of adaptive and neutral processes in explaining observed levels of genetic variation [4]. However, adaptation does not occur in isolation, but acts on genetic variation that is also shaped by mutation, recombination, gene flow, and genetic drift [5]. More recently there has been increased effort in understanding how these fundamental evolutionary forces operate in concert to generate and maintain the levels of genetic diversity commonly observed in natural populations [6,7].

The increasing feasibility of high-throughput sequencing and genotyping, alongside subsequent characterisation of genome-wide variation across large numbers of individuals, has revealed that at the genomic level, patterns of variation and divergence among natural populations and species are highly heterogeneous [8]. A key feature of these “genomic landscapes” of divergence that has received particular attention is the presence of so-called “islands of differentiation”: outlier regions of the genome with high levels of divergence estimated from statistics such as *F*_*ST*_ or *d*_*xy*_ that consistently emerge at the same loci in different comparisons between populations or related species [8–11]. Initially these regions were termed “islands of speciation”, and were thought to arise as a result of reduced gene flow in genomic regions associated with reproductive isolation [8,12]. Subsequent research has revealed that highly heterogeneous patterns of genomic divergence can occur even in the complete absence of gene flow, as a result of recombination rate variation and linked selection [13,14]. In genomic regions of low recombination, selection for beneficial mutations (positive selection), or against deleterious mutations (background selection), will impact relatively large genomic regions as a result of high levels of linkage disequilibrium (LD) among sites. Selection within these regions reduces diversity within populations, and increases levels of differentiation among them, resulting in “islands” of increased differentiation that persist over evolutionary time [14,15]. Another, less well explored reason for which islands of divergence can arise is due to the differential effects of genetic drift in response to variation in effective population size across different genomic regions; something that may be particularly important in recently colonised populations [16,17]. These circumstances promote fixation of haplotypes and therefore result in either reduced or inflated local differentiation.

Comparing patterns of genomic differentiation among sets of populations or species at different stages of the divergence/speciation continuum is a powerful way of disentangling the forces that shape variation among populations. Martin et al. [18] showed that, across multiple *Heliconius* butterfly populations and species, patterns of genomic variation were shaped by a combination of gene flow and selection, particularly in genomic regions harbouring genes involved in wing patterning. In contrast, Renaut et al. [10] showed that in *Helianthus* sunflowers, genomic architecture was the main driver of genomic differentiation across sets of populations. Similarly, recent research in birds has revealed that differentiation landscapes are conserved across populations, species and even across avian families, with the same islands of differentiation arising among populations of distantly related species [19–21]. This latter pattern appears to have arisen, at least in part, as a result of a highly conserved synteny and recombination landscape in birds [22], with background selection in regions of low recombination producing recurrent islands of differentiation [23].

It is now clear that the recombination landscape and linked selection are key drivers of genomic variation within and among populations. However, we are only just beginning to understand how this linked selection interacts with other evolutionary forces to shape patterns of differentiation across natural populations and species [23–28]. A recent, large-scale analysis of threespine sticklebacks (*Gasterosteus aculeatus*) showed that islands of differentiation were more likely to arise in low recombination regions when gene flow occurred between populations [29]. There is also a significant impact of divergence time; in recently separated populations the differentiation landscape is most likely to reflect selective sweeps. Then, as divergence accumulates, genomic architecture is expected to play an increasingly important role in generating these genomic islands [23].

Widespread continental species are excellent models for studying how demography and the environment shape genetic and phenotypic variation among populations, due to their large effective population sizes and ecologically varied ranges. Insight into the evolutionary history of such species can be gained if genetic variation is characterized across much of its geographical range. Cross-population comparisons of genetic variation can then be utilised to make inferences about phylogeography, levels of gene flow between populations and how adaptation to different environmental and ecological conditions occurs [30]. The first large-scale studies were performed in humans - i.e. the HapMap Projects [31–33] which characterized human genetic variation on different continents, with a view to determining the feasibility of association mapping studies. Similar studies have been conducted in domesticated species and their wild ancestors [34–36], and in model organisms [37,38]. More recently, there is a growing appreciation that HapMap-type studies are useful for studying signatures of selection and adaptation in natural populations of species with large effective population sizes and high levels of gene flow [39–42].

The European great tit (*Parus major major*) is an excellent model for ecological and evolutionary studies [43]. As is the case with several avian species which are amenable to long-term study [44], a wealth of ecological data exists across multiple great tit populations [45–47], enabling informed hypotheses about selection to be tested in this system. Phyleogeographic research using mitochondrial DNA suggests that this species has recently expanded across its European range, possibly from a single refugium in South-East Europe [48]. Contemporary populations are characterised by large effective population sizes and low levels of genetic differentiation [49,50]. However, these previous cross-population molecular studies have relied on a modest number of microsatellite loci and mitochondrial DNA, making the detection of genomic regions under selection impossible. The genome of the great tit has recently been sequenced [51], and a high density panel of *ca* 500,000 SNP markers has been developed [52]. A recent study of two European populations using this marker panel suggests that rapid adaptation has occurred at the genomic and phenotypic levels, with pronounced selection on morphology [53].

Here, we perform a HapMap study of 647 unrelated individuals across 29 populations (Fig. 1), to examine how genomic architecture, natural selection and population history have shaped patterns of genomic variation across recently colonised European great tit populations. Using a large SNP panel typed across all individuals, we first characterise genome-wide patterns of variation within and among populations, in order to infer population history. We then examine how variation is partitioned across the genome, and test the hypothesis that highly divergent genomic regions have arisen in genomic regions of low recombination [13,14]. Finally, we examine how genomic divergence accumulates along the colonisation route of this species, with the aim of inferring how recent natural selection and demography drive variation across the genome in the wild.

**Figure 1.**
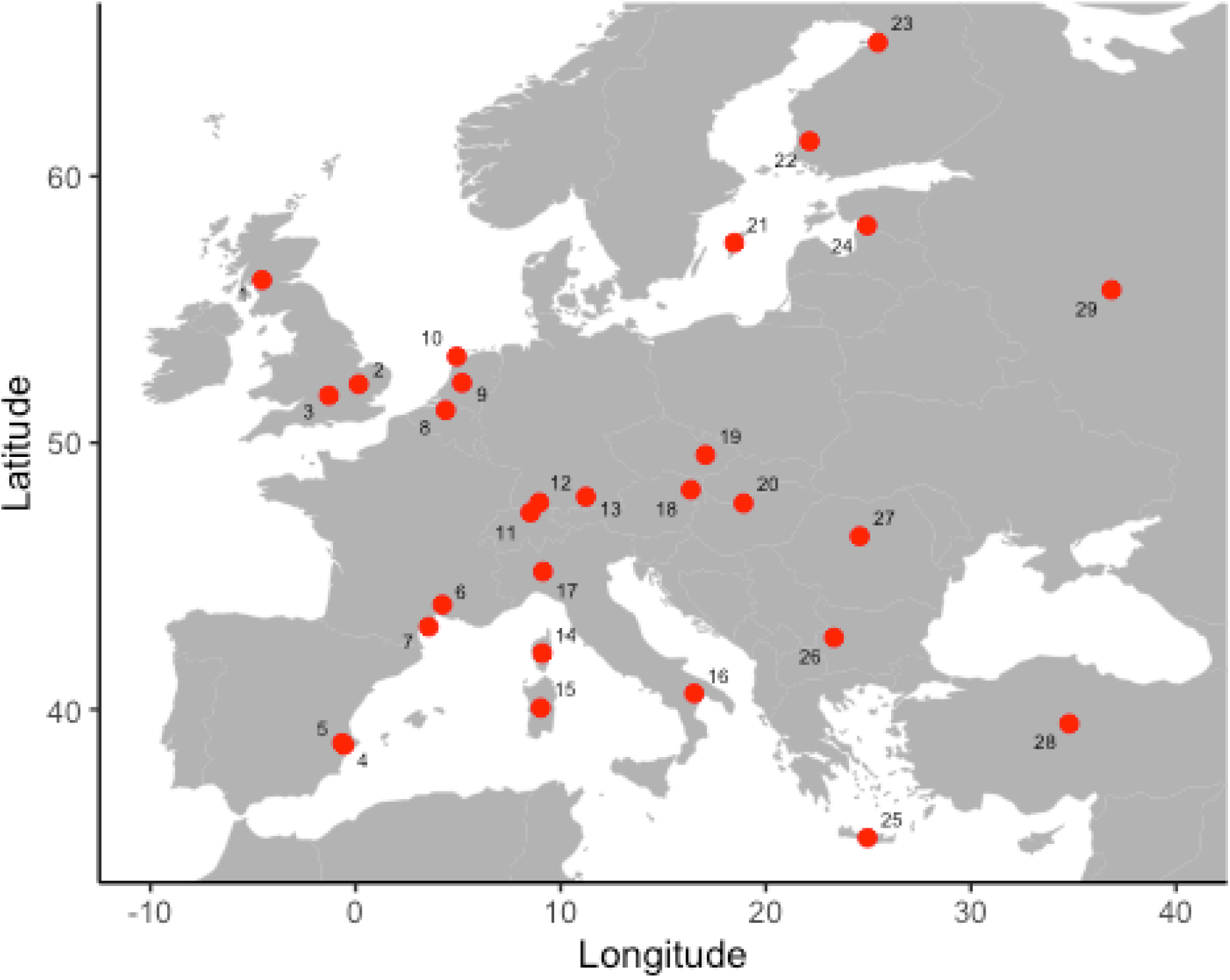
Sampling locations of great tit populations. Population names and sample sizes are given in Table S1, and numbers on the map correspond to the “code” column in Table S1.

## Results and Discussion

### Genetic diversity and population history

Sampling locations and sample sizes for each population are given in Table S1. Levels of genetic diversity *(π*_*SNP*_) were generally high, but we observed substantial differences among populations (Fig. 2A). Similarly, LD declined rapidly with genomic distance in all populations, reaching baseline levels within ∼5kb in all populations, but also varied among populations (Fig. S1). Highest levels of LD (and lowest levels of genetic diversity) were observed in the Mediterranean island populations of Crete (Greece) and Sardinia (Italy), with lowest levels of LD in central and western Europe (Fig. S1). This is consistent with reduced effective population size in these island populations, either as result of the colonisation process or more recent bottlenecks, along with low levels of subsequent gene flow from the continent to the islands [54,55].

**Figure 2.**
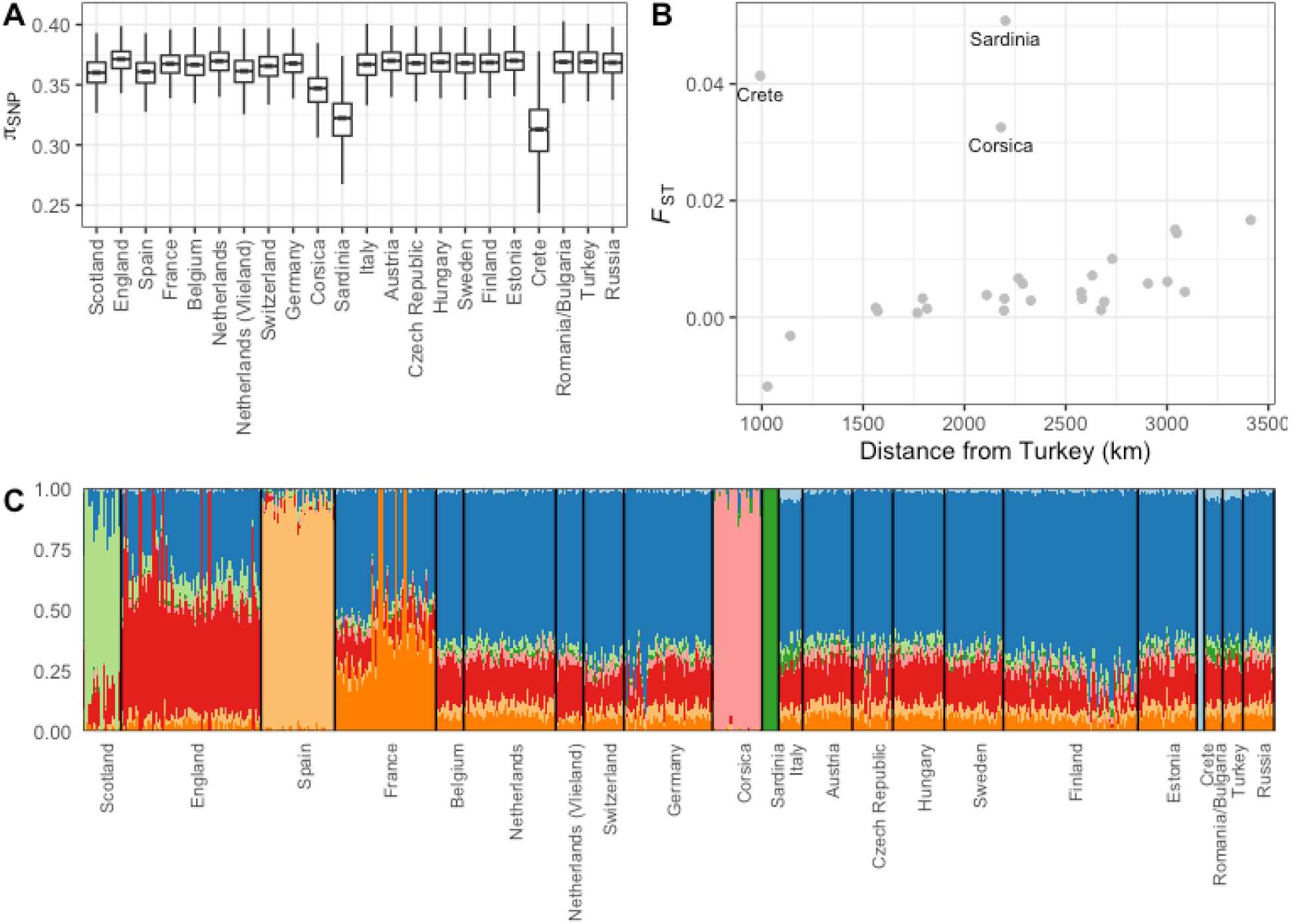
Genetic diversity and structure in European Great tit populations. **A** Nucleotide diversity within each population. **B** Pairwise *F*_*ST*_ in relation to geographic distance from the Turkey, only including comparisons involving Turkey. **C** Output from Admixture analysis at *K* = 8. Population details can be found in Table S1.

Mean genome-wide *F*_*ST*_ between all pairs of European great tit populations was 0.015, with no significant pattern of isolation-by-distance (Mantel test; r = 0.13, p = 0.18; Fig. S2). Instead, the highest levels of *F*_*ST*_ were found in comparisons involving the Mediterranean island populations of Corsica (France), Sardinia and Crete (Fig. S2). Admixture analysis was consistent with this pattern (Fig. S3); the *K* = 2 analysis assigned individuals in Sardinia and Corsica to one genetic cluster, and the remaining populations to the second. Thus, it is likely that much of the genetic structure between European great tit populations is a result of genetic drift in these small island populations. Admixture analysis also revealed some structure between (mainly peripheral) mainland and larger island populations. At *K* = 3 (the model that best fitted the great tit data; Fig. S4), Spain was separated from the rest of mainland Europe. Increasing values of *K* resulted in the separation of populations in Scotland (*K* = 4), Sardinia (from Corsica; *K* = 5), southern France (*K* = 6), Crete (*K* = 7) and England (*K* = 8). The Admixture output at *K* = 8 is displayed in Fig. 2C as this gives the most detailed picture of genetic structure among European great tit populations. Further increases in *K* did not generate patterns of structure that corresponded to geographical variation (Fig. S3), and were increasingly less well supported (Fig. S4). Thus, even with hundreds of thousands of markers Admixture was unable to separate many of the European populations, confirming that levels of divergence are extremely low [51]. PCA largely corroborated the Admixture results, with PC1 separating Corsica and Sardinia from the remaining populations, PC2 separating Spain, while PC3 and PC4 separated Scotland, England, Corsica, Sardinia and Crete (Fig. S5).

Maximum likelihood analyses implemented in TreeMix showed that a model with no migration explained 97.8% of variance in relatedness between populations [56]; increasing the number of migration events substantially improved the percentage of relatedness explained, up to 99.7% when 10 migration events were fitted (Fig. S6). In Figure 3 we display the maximum likelihood tree with three migration events, after which the variance in relatedness explained plateaued when more migration events were added (Fig. S6). The tree was generally characterised by short branch lengths, with the exception of the island populations of Sardinia and Crete, which were grouped with the population from mainland Italy (Fig. 3). Thus, the TreeMix analysis is consistent with large populations and low overall genomic divergence, with the exception of the Mediterranean island populations. However, much (though not all) of the grouping that did occur among continental populations made geographical sense, with populations from Finland and Estonia grouped together, as were some populations from South-East Europe, and populations from England and Scotland (Fig. 3). Interestingly, TreeMix grouped the Spanish and Corsican populations, which is consistent with previous subspecies descriptions of European great tits [57]. The three fitted migration edges all involved Sardinia, with migration from eastern Europe to Sardinia, and from Sardinia to Corsica and Spain (Fig. 3).

**Figure 3.**
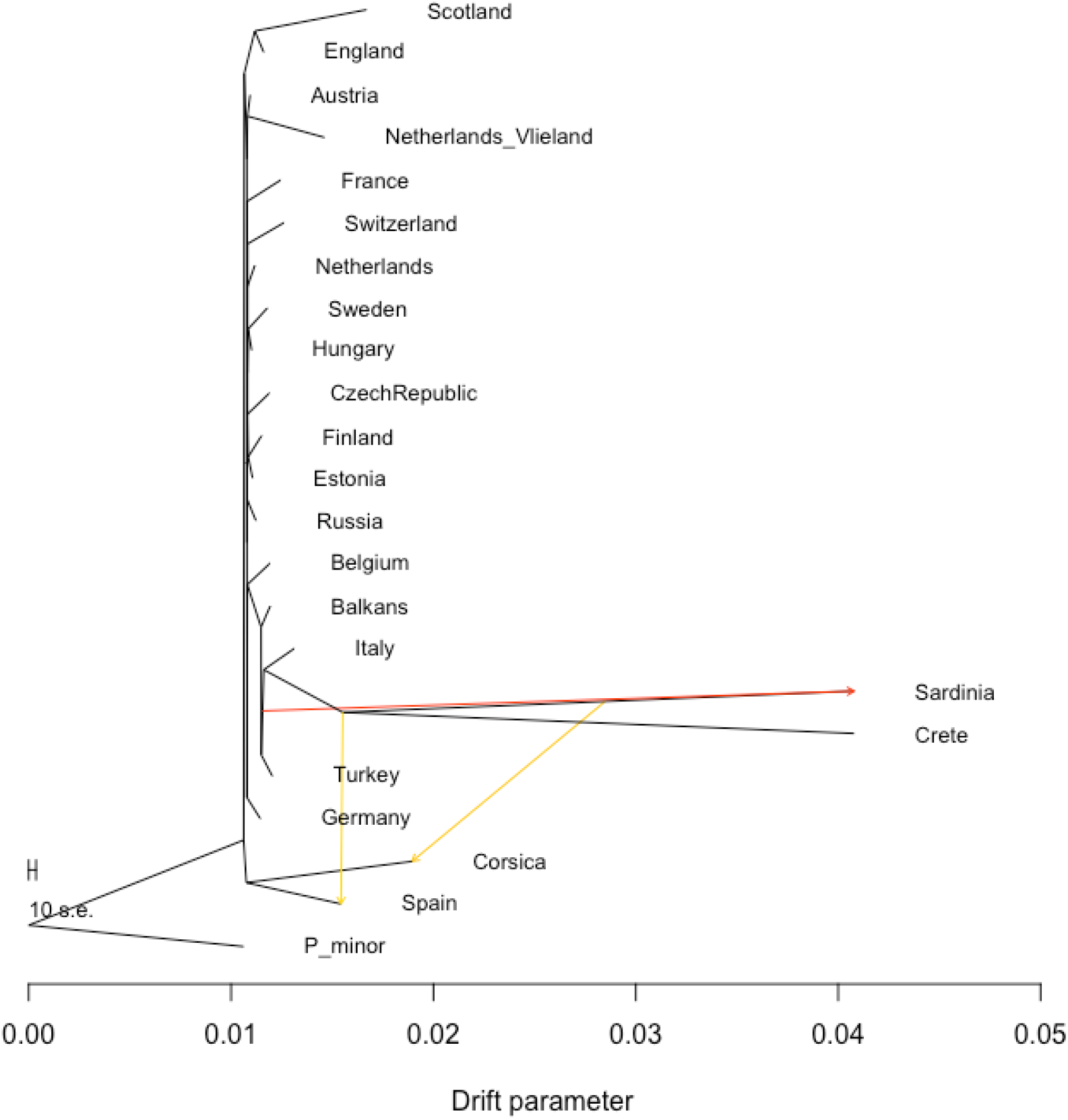
Maximum likelihood tree inferred by TreeMix, allowing three migration events. The two migration events (arrows) are coloured according to their weight (red = higher migration), and horizontal branch lengths are proportional to the amount of genetic drift that has occurred along the branch. A population of the great tit’s sister species, *Parus minor*, was used as an outgroup. Population details are given in Table S1.

We next tested the hypothesis that great tits colonised Europe from a single refugium in South-East Europe. This scenario has been suggested before [48], but due to the low number of genetic markers available there has been limited power with which to test this hypothesis. Using our genome-wide panel of SNP markers, we compared genetic and geographic distance between each population and the proposed refugial populations. Because of the elevated structure in Corsica, Sardinia and Crete (Fig. S2), we excluded comparisons involving these populations. We found that *F*_*ST*_ was significantly related to distance from Turkey (r = 0.81, p < 0.001; Fig. 2B) and Romania/Bulgaria (r = 0.44, p = 0.001). The same relationship was not found for alternative potential refugial populations [58] in Spain (r = −0.09, p = 0.55), or southern Italy (r = 0.04, p = 0.77). Our results therefore lend empirical support to the hypothesis [48] that great tits colonised Europe primarily from a single refugium in the south-east. Clearly, although our sampling was extensive, it is not exhaustive, and more fine-scaled sampling in eastern Europe would be required to determine the extact location and extent of refugial great tit populations. Sampling in North Africa would also be useful to determine whether further refugia exist, and to quantify the extent of admixture between European and African great tit populations.

### Genomic landscapes of differentiation

It is likely that many, and perhaps the majority, of wild populations are characterised by highly heterogeneous patterns of differentiation across the genome [27]. This is the case in European great tits - despite the extremely low average *F*_*ST*_, we found genomic regions with very high levels of genomic structure (maximum *F*_*ST*_ for 10kb and 500kb windows was 0.98 and 0.07, respectively). To examine how landscapes of genomic divergence have formed along the colonisation route of European great tits, we calculated windowed *F*_*ST*_ in 500kb bins between each population and the proposed refugial population in Turkey. We found that *F*_*ST*_ varied markedly across the genome in all comparisons (Fig. 4; Fig. S7). Outlier regions (windows with standardized *F*_*ST*_, hereafter *zF*_*ST*_, > 10) were found in all comparisons apart from Crete and Sardinia, in which overall levels of divergence were highest, with some outlier regions found across multiple comparisons (Fig. 4). Our results suggest, therefore, that genomic islands of differentiation can and do arise even among recently separated populations.

**Figure 4.**
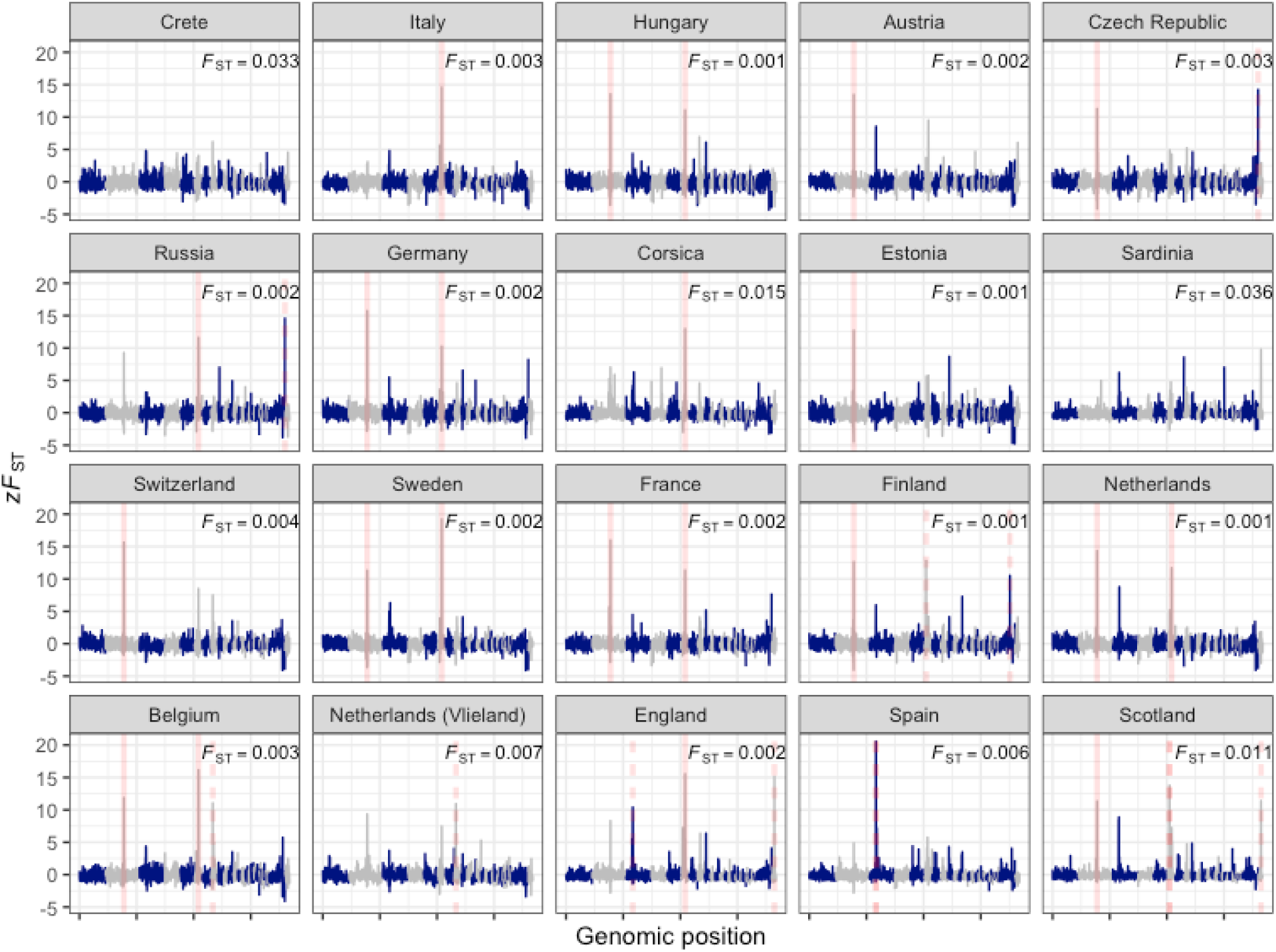
Landscapes of relative genomic differentiation in European great tit populations. *zF*_*ST*_ across the genome is averaged in 500kb windows, with each panel displaying a pairwise comparison with the proposed refugial population in Turkey. Red lines represent *F*_*ST*_ outliers (windows with mean *F*_*ST*_ values at least 10 standard deviations greater than the global mean for that comparison) shared across more than two comparisons (solid red lines), or specific to one or two comparisons (dashed red lines). Mean, untransformed *F*_*ST*_ values are given in the top-right of each panel, and are fully displayed in Fig. S7.

Considering all populations, *F*_*ST*_ calculated in 500kb windows was strongly negatively correlated with local variation in recombination rate (Spearman correlation, r = −0.50, p < 0.001). Although recombination rate and gene density were positively correlated, (r = 0.16, p < 0.001), *F*_*ST*_ was only moderately correlated with gene density in 500kb windows (r = −0.09, p < 0.001), and this correlation became weak when calculated in 10kb windows (r = −0.01, p < 0.001). Thus it appears that the negative relationship between *F*_*ST*_ and recombination rate is not driven by gene density. Examining how the relationship between genomic differentiation and recombination varied among populations revealed that *F*_*ST*_ was negatively related to recombination rate in almost all comparisons with Turkey (Fig. 5). The relationship between *F*_*ST*_ and recombination rate was generally weak, but in a handful of populations this relationship was substantially stronger - most notably in the island populations of Corsica, Sardinia and Crete, and in the peripheral mainland populations of England, Scotland, Spain, Finland and France, which are the populations that are likely most susceptible to drift (Fig. 5).

**Figure 5.**
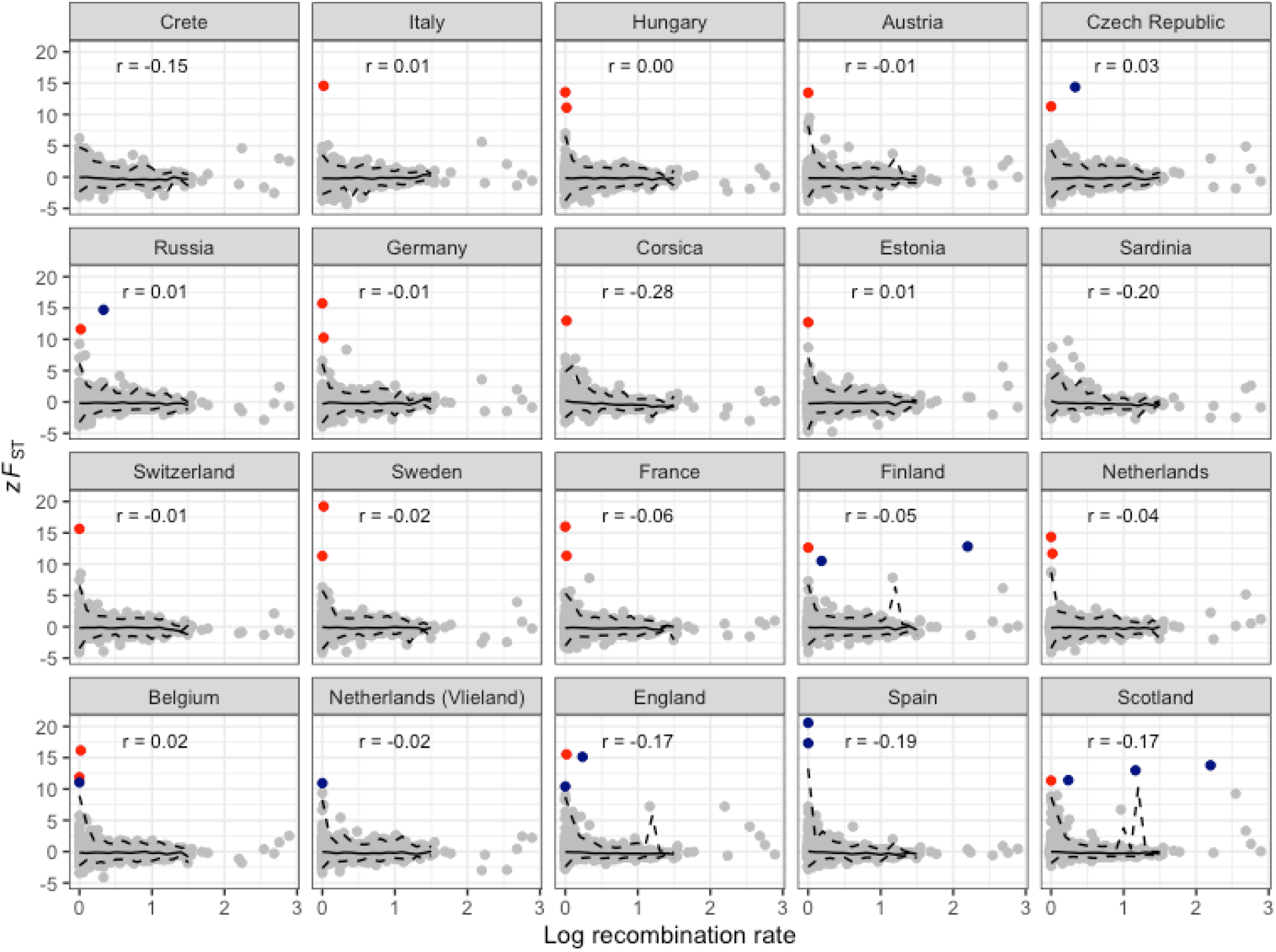
Genomic differentiation and recombination rate variation in European great tit populations. Each point is the mean of a 500kb window, with each panel displaying a pairwise comparison with the proposed refugial population in Turkey. Coloured points represent *F*_*ST*_ outliers (mean standardized *F*_*ST*_ values of *zF*_*ST*_ > 10) shared across more than two comparisons (red), or specific to one or two comparisons (dark blue). Solid and dotted lines represent median and 99% quantiles of *F*_*ST*_ windows from bins of 0.1 log cM/Mbp.

As a further examination of how allele frequency variation and haplotype structure may be shaped by neutral and adaptive processes, we compared *F*_*ST*_ and recombination rate with *Rsb* - a measure which aims to detect regions of the genome under positive selection by comparing extended haplotype homozygosity profiles between populations, and is expected to be less sensitive to variation in recombination rate because extended haplotypes should be present in both populations when recombination frequency is lower [59]. Specifically, we compared the distributions of these statistics in 500kb windows using three populations in the Netherlands, Finland and Spain, with the aim of exploring how *Rsb* varies with recombination rate and with *F*_*ST*_. We found that, across the genome, correlations between absolute *Rsb* and recombination rate were only slightly weaker (Spearman rank r between −0.25 and −0.20) compared to those between *F*_*ST*_ and recombination for the same pairwise comparisons (r between −0.36 and −0.20), and that the regions of high *F*_*ST*_ in these comparisons tended to be in these same regions of low recombination and high *Rsb* (Fig. S8). Notably, though, there were several cases where windows with extreme *Rsb* values occurred in regions of high recombination, and most high *F*_*ST*_ regions in high recombination regions were also *Rsb* outliers (Fig. S8). Below we explore the possibility that such regions are strong candidates for recent adaptive evolution.

Background selection is a key driver of genomic differentiation in birds [20,21], and other organisms [60], but is not expected to play a major role in driving islands of differentiation over the timescales (tens of thousands of years) relevant to this study [23,29]. On the other hand, differential effects of drift operating across the genome could contribute substantially to the highly heterogeneous patterns of differentiation observed in European great tits. Our finding that the relationship between recombination rate and *F*_*ST*_ is strongest in populations experiencing the greatest levels of drift (e.g. islands; Fig. 5) is consistent with this - low recombining regions have reduced effective population size and increased rates of lineage sorting due to drift. Drift has been suggested to be a driver of heterogeneous genomic landscapes in other systems [16,17], and our study suggests that it may play a key role in shaping genomic structure in widespread, continental species.

Outlier regions of very high differentiation (*zF*_*ST*_ > 10) mostly occurred in areas of the genome with low recombination rates (Fig. 5) and the recombination rate of outlier regions was (just) significantly lower than the genome-wide average (Wilcoxon test, *P* = 0.048). Of the 11 outlier regions, nine were found in only one or two comparisons, while the other two were found in 12 and 10 comparisons, respectively (Table S2). We hereafter refer to outlier regions found in one or two comparisons as “population-specific” outlier regions, and to those found in more than two comparisons as “shared” regions.

Regions of high differentiation that are not shared among populations could potentially arise as a result of recent drift, but are also candidate regions for recent positive selection [23]. Several lines of evidence suggest that this may be the case in European great tits. Firstly, population-specific regions tended to occur in regions of higher recombination than shared outliers, although low sample size prevents a formal statistical analysis of this. Second, population-specific outliers tended to be found in the most peripheral European great tit populations, with three found in Scotland, and two in England, Spain and Finland; the remaining outlier regions were found in comparisons involving the Czech republic, Russia, Vlieland (Netherlands) and Belgium (Table S2). Notably, these populations are not the lowest in genetic diversity and therefore drift effects are not necessarily stronger in these peripheral populations. Third, the *Rsb* tests, which indicate which population has undergone a selective sweep, also tend to find evidence of positive selection in the more peripheral population, as expected under an adaptive evolution scenario (Fig. S8). This directionality cannot be assessed with *F*_*ST*_, because it is a combined measure of allele frequency differences. Of course, across the genome, both positive selection and linked background selection are likely to be operating. Observational and experimental research shows that adaptation at range edges is a key feature shaping divergence among recently colonised and expanding populations [61–63]. Population-specific regions appeared less likely to be in regions of low recombination than shared regions (Fig. 5), although the small number of regions precluded testing this hypothesis formally. Thus, it is likely that genomic architecture plays a key role in determining how both selection and drift have shaped genomic variation across the recent evolutionary history of European great tits.

To further explore how selection may have shaped variation in *F*_*ST*_ outlier regions, we estimated levels of nucleotide and haplotype diversity within these regions. Nucleotide diversity (π_*SNP*_) in outlier regions varied from 0.21 to 0.48, and diversity in these regions was significantly lower than the genome-wide average (Wilcoxon test, *P* = 0.018; Fig. S9A). However, there appeared to be little difference in nucleotide diversity between shared and population-specific regions (Fig. S9A). Haplotype diversity varied substantially among regions, with haplotype richness ranging from 72 to 1033. Both haplotype richness and marker density in shared regions tended to be lower than those in population-specific regions (Fig. S9B), consistent with lower recombination frequency. A detailed examination of haplotype structure in one shared and one population-specific region is displayed in Figure S10. The population-specific outlier region (to Finland, situated on chromosome 1A) was characterised by a complex structure, with a single haplotype at high frequency in Finland compared to other populations, indicating a population-specific selective sweep and high background diversity (Fig. S10). In contrast, the shared region on chromosome 2 was much less complex, demonstrating high frequency haplotypes that are found across a range of populations. Our data therefore suggest that examining patterns of haplotype diversity in outlier regions may help to separate recent episodes of positive selection from drift and background selection (Figs S9, S10).

Potential candidate genes found within shared and population-specific outlier regions are displayed in Table S2. Perhaps most notable among these is *COL4A5*, a gene found to be associated with bill length, and under selection between populations in England and the Netherlands, in a recent great tit study [53]. Here we found that the *COL4A5* region is an *F*_*ST*_ outlier in England and Scotland, but not in any other European populations (Table S2). UK great tits have been described as a separate subspecies based on beak shape [64], and our results here, combined with previous results, suggest that this divergence is the result of recent natural selection in the UK [53]. Another notable candidate gene potentially involved in beak morphology, and previously found to be under selection in UK great tits is *BMPR1A*, which plays a key role in palate development [65] and in this study was found in an outlier region in Scotland. Other candidate morphology and obesity genes in the population-specific outlier regions in the UK included *PPP1CB*, which may play a role in adipogenesis [66] and *GHITM*, which appears to have been subject to natural selection in human pygmy populations [67]. Thus, morphological traits may frequently be involved in adaptation in great tits.

In addition to morphological candidates in the UK, we found outlier regions specific to cold populations in Scotland, Finland and Russia (Table S2), containing at least one candidate gene for thermal stress (*CDKN1B*) [68]. Other genomic outlier regions contained potential candidate genes associated with malaria infection (*MRPL33*) [69] and colour variation (*SOX10*) [70]. This is thus far an exploratory analysis, and we are therefore reluctant to speculate whether these candidate genes are genuine targets for natural selection, and more reluctant still to speculate as to how selection might be driving variation at these regions. Regardless, these candidates will provide useful starting points for future genomic and ecological investigation.

HapMap style projects have been hugely informative in shaping our understanding of how natural selection operates in humans and other model species [31,38]. This study is one of the largest to date of genomic variation in a wild vertebrate, which has helped to reveal the evolutionary history of great tits, and to identify candidate genes and traits that may have been involved in adaptation during and/or after postglacial recolonisation. Further, this work will form the foundation of many future analyses. Clearly, we have only touched on haplotype-based methods to infer adaptation here, and this will be the subject of future work. Environmental association approaches are also highly suited to detecting adaptation in widespread continental species [71,72], and further work will test how variation in the environment has shaped patterns of genomic variation in great tits [73]. This combination of environmental and genomic data in species such as great tits, in which a wealth of ecological and genomic resources are available, is likely to generate interesting insights into the the genetic and phenotypic basis of natural selection.

## Materials and Methods

### Sampling and molecular methods

Samples were collected from 29 populations from 22 regions across Europe (Fig. 1; Table S1). Samples were pooled into regions either based on geographical proximity (e.g. Cambridge and Wytham woods), or based on sample size (e.g. Romania and Bulgaria). An exploratory analysis considering all sampled populations separately yielded virtually identical results to those shown here, and in no cases did we observe substructure within pooled populations in our Admixture analyses (Fig. S3).

Birds were trapped from nest boxes, or using mist nets, and ringed with a uniquely numbered aluminium ring. Blood was taken via brachial or tarsal venipuncture, and stored in either 1 ml Cell Lysis Solution (Gentra Puregene Kit, Qiagen, USA), Queen’s buffer, or absolute ethanol. All samples were genotyped using a custom-made Affymetrix® great tit 650K SNP chip at Edinburgh Genomics (Edinburgh, United Kingdom), following the approaches outlined in [52]

### Analyses

Unless stated otherwise, all population genetic statistics were calculated in PLINK version 1.9 [74], and downstream analysis and plotting was carried out in R version 3.3 [75]. For population genetic analyses, we used the filtering approaches outlined in [53]. Briefly, we randomly removed individuals from pairs with relatedness values > 0.4, and for demographic analyses we used a LD pruned dataset (based on VIF > 0.2), with SNPs associated with an inversion on chromosome 1A [**???**] removed. After filtering, a total of 647 samples typed at 483888 SNPs were retained for analysis.

In each population, we estimated LD (*R*^2^) for each pair of markers within 50kb on the same chromosome, and compared this to physical distance between marker pairs. We calculated observed heterozygosity for each SNP and population using a reduced SNP dataset, which was pruned based on LD to remove all markers with *R*^2^ > 0.1, then thinned with a probability of retaining each variant of 0.25. We calculated genome-wide (mean) *FST* between each pair of populations using the pruned and thinned dataset described above. Pairwise *F*_*ST*_ was compared to geographic distance between populations using Mantel tests, implemented in the Ecodist package in R [76]. We tested whether genetic structure was related to distance from candidate refugial populations (in Romania/Bulgaria, Turkey, Spain and Italy), using Pearson correlations. We also estimated population structure using Admixture version 1.3, with default settings [77]. We varied values of *K* from one to ten; by which point increasing values of *K* provided no informative information about population structure (see results). Model support for each value of *K* was estimated by calculating 5-fold cross-validation error. Finally, we visualised the evolutionary history among European great tit populations by generating a maximum likelihood tree in TreeMix version 1.13 [56]. We rooted the tree using a sample of *P. minor* individuals sampled from Amur, Russia [52]. We fitted models allowing for range of migration events (0-10), and used a window size of 500 SNPs [56]. To assess model fit, we calculated the proportion of variance in relatedness between populations explained by each model [56].

Recombination rates at each locus were estimated by comparing the location of SNPs on the genome assembly (v1.1) with their location on the great tit linkage map [78]. Previous linkage mapping, using a lower density SNP chip, was independently carried out in UK and Netherlands great tit populations and the two maps were almost identical [78]. For the purposes of this analysis, we used SNPs and marker intervals from the UK comprehensive map. A total of 2706 SNPs were located on both the genome assembly and the linkage map. Thus a positon in Mb and cM of each of these SNPs is known. All other SNPs on the HD chip have a physical (Mb) position but no known linkage map position. The great tit genome v1.1 is 1.02 Gb long, so the average physical interval between mapped SNPs is ∼376Kb. The linkage map position of each unmapped SNP was estimated by interpolation; by taking the closest mapped SNP in either direction, and assuming a constant recombination rate in the interval between those SNPs. For example, an unmapped SNP with physical position 1.4Mb, flanked by mapped SNPs at 1.0Mb/0.5cM and 2.0Mb/1.0cM, would be estimated to be located at 0.5 + (1.4-1.0)/(2.0-1.0) * (1.0-0.5) = 0.7cM. Having interpolated cM position of every SNP, the local recombination rate was calculated as the cM interval spanned by the nearest neighbouring SNPs, divided by the physical distance (bp) spanned by those same neighbouring SNPs. In other words, for the ith SNP, the recombination rate is estimated as the linkage distance between the i-1th and i+1th SNP, divided by the physical distance between the i-1th and i+1th SNP. For downstream analyses, local recombination rates were estimated by averaging across all SNPs in each 500kb window. We calculated gene density in 10kb and 500kb windows using custom R scripts and the annotated great tit genome (v 1.1) [51].

We examined the genomic landscape of differentiation across European great tit populations by calculating *F*_*ST*_ in 10kb and 500kb bins, using python scripts obtained from Github (https://github.com/simonhmartin/genomics_general). We did not estimate *d*_*xy*_, as this parameter is difficult to estimate accurately from single SNP loci [14]. We also calculated standardised *F*_*ST*_ (*zF*_*ST*_) by mean-centring windowed values and dividing them by the standard deviation among windows. We defined outlier regions as 500kb bins with *zF*_*ST*_ values greater than ten. Finally, *Rsb* [59] was calculated for three pairwise comparisons (in Spain, Finland and the Netherlands), using the R package Rehh [79], and this averaged into 500kb windows. Unlike *F*_*ST*_, the *Rsb* statistic gives an indication of which population an adaptively important haplotype is under positive selection in, and is thought to be less sensitive to local recombination rates [59]. *Rsb* estimation requires phased genotype data, so phasing was performed using shapeIT2 v 2.r837 [80]. The –duohmm argument was used to ensure that family information, where available, was used to improve the accuracy of phasing. The –effective-size parameter was set at 500,000 reflecting the large effective population size of European great tit populations [51]. Local recombination rates (measured in cM/Mbp) were used in the map files. Comparisons between *F*_*ST*_, *Rsb*, recombination rate and gene density were carried out using Spearman rank correlation (as all statistics were highly skewed).

### Ethics Statement

Great tit DNA samples were provided by researchers studying hole-nesting great tit populations with permissions and licenses to study and sample birds granted by the following organisations: Bundesministerium für Wissenschaft und Forschung (Ref. II/3b Gentechnik und Tierversuche), Czech Republic Ministry of Education Youth and Sports (MSMT-56147/2012-310), Dutch legal entity Dier Experimenten Comissie (NIOO 14.11, NIOO 07.04), Dutch Central Authority for Scientific Procedures on Animals (AVD-801002017831), Estonian Ministry of Agriculture - Animal Procedures Committee (License 108), Ethical Committee of the Canton of Bern, Ethics Committee for animal experimentation of Languedoc Roussillon CEEA-LR (APAFIS#8608-2017012011062214), Ethics Committee of Lomonosov Moscow State University (Approval numbers 26ch, 37zh, 49zh, 59zh, 88zh), Federal Office for the Environment - Switzerland (License 234), Flemish Ministry - Agency for Nature and Forest, Generalitat Valenciana - Conselleria de Medi Ambient, Hellenic Ministry of Environment and Energy, Istituto Superiore per la Protezione e la Ricerca Ambientale - ISPRA (Law 157/1992 Art. 4 (1) and Art. 7 (5)), Mehmet Akif Ersoy University - Local Ethical Committee on Animal Experiments (93773921-9), Middle Danube Valley Inspectorate for Environmental Protection, Nature Conservation and Water Management (43355-1/2008), Regierung Oberbayern (55.2-1-54-2532-140-11), Regierungspräsidium Freiburg - Departments for Animal Welfare and Nature Conservation, Regional State Administrative Agency of Southern Finland (ESAVI/1650/04.10.03/2012, ESAVI-1804/04.10.03/2011), Romanian Academy of Sciences (Permit number 2257), Swedish Committee for Experiments on Animals (Licence number C 108/7), UK Home Office (License PPL 30/2409), University of Oxford - Department of Zoology Local Ethical Review Committee, University of Glasgow AWERB committee, Veterinary Office of the Canton of Zurich (License 195/2010) Samples were imported to the UK under DEFRA general import license IMP/GEN/2011/01, according to regulations outlined by the UK Animal Health and Veterinary Laboratories Agency.

## Acknowledgements

This work was supported by grants from the Natural Environment Research Council (grant NE/J012599/1 to J.S. and B.C.S) and the European Research Council (grant 202487 to J.S. and grant 339092 – E-Response to MEV) L.G.S was supported by fellowships from the Edward Grey Institute for Ornithology and the BBSRC (BB/N011759/1). We thank Claire Bloor, Geoff Scopes and Alessandro Davassi of Affymetrix for their help during the chip design and genotyping calling processes. Richard Talbot and Alison Downing of Edinburgh Genomics provided the genotyping service.

## Data Availability

All data and code to reproduce the results are available on Github: https://github.com/lgs85/SpurginBosse_Hapmap

