## Supplementary Material for "The great tit HapMap project: a continental-scale analysis of genomic variation in a songbird"

**Table S1** Sampling details for the 29 European great tit populations. “Code” corresponds to labels in Figure 1 of the main text. Sample size (N) is after pruning (see Methods).

| Code | Population | Region | N | Latitude | Longitude |
|---:|---|---|---:|---:|---:|
| 1 | Loch_Lomond_Scotland | Scotland | 21 | 56.108 | -4.562 |
| 2 | Cambridge_UK | England | 28 | 52.198 | 0.152 |
| 3 | Wytham_UK | England | 48 | 51.776 | -1.303 |
| 4 | Font_Roja_Spain | Spain | 19 | 38.664 | -0.544 |
| 5 | Mariola_Spain | Spain | 21 | 38.737 | -0.658 |
| 6 | La_Rouviere_France | France | 19 | 43.930 | 4.235 |
| 7 | Montpellier_France | France | 36 | 43.098 | 3.569 |
| 8 | Antwerp_Belgium | Belgium | 15 | 51.226 | 4.390 |
| 9 | Westerheide | Netherlands | 50 | 52.250 | 5.190 |
| 10 | Vlieland_NL | Netherlands (Vlieland) | 15 | 53.238 | 4.932 |
| 11 | Zurich_Switzerland | Switzerland | 22 | 47.383 | 8.523 |
| 12 | Radolfzell_Germany | Germany | 18 | 47.759 | 8.957 |

| Code | Population | Region | N | Latitude | Longitude |
|---|---|---|---|---|---|
| 13 | Seewisen_Germany | Germany | 30 | 47.978 | 11.233 |
| 14 | Pirio_Muro_Corsica | Corsica | 27 | 42.110 | 9.104 |
| 15 | Sardinia | Sardinia | 9 | 40.056 | 9.025 |
| 16 | Matera_Italy | Italy | 5 | 40.601 | 16.514 |
| 17 | Po_Plain_Italy | Italy | 8 | 45.171 | 9.138 |
| 18 | Vienna_Austria | Austria | 27 | 48.239 | 16.347 |
| 19 | Velky_Kosir_Czech_Republic | Czech Republic | 22 | 49.547 | 17.062 |
| 20 | Pilis_Mountains_Hungary | Hungary | 28 | 47.733 | 18.917 |
| 21 | Gotland_Sweden | Sweden | 32 | 57.504 | 18.466 |
| 22 | Harjavalta_Finland | Finland | 44 | 61.314 | 22.139 |
| 23 | Oulu_Finland | Finland | 29 | 65.013 | 25.468 |
| 24 | Kilingi-Nõmme_Estonia | Estonia | 32 | 58.150 | 24.938 |
| 25 | Crete | Crete | 4 | 35.183 | 24.960 |
| 26 | Bulgaria | Romania/Bulgaria | 3 | 42.696 | 23.328 |
| 27 | Romania | Romania/Bulgaria | 7 | 46.497 | 24.582 |
| 28 | Turkey | Turkey | 11 | 39.464 | 34.797 |
| 29 | Zvenigorod_Russia | Russia | 17 | 55.738 | 36.857 |

**Table S2** Details of outlier $F_{ST}$ regions in European great tit populations. For each region, $F_{ST}$ was calculated for each population versus Turkey. The "number of outliers" represents the number of *populations* in which each region was detected as an outlier. Selected candidate genes found within 500kb outlier windows (see main text) are displayed.

| Chromosome | Position (MB) | Number of outliers | Outlier comparisons | Candidate genes |
|---|---|---|---|---|
| 2 | 81.5 | 12 | Austria, Belgium, Czech Republic, Estonia, Finland, France, Germany, Hungary, Netherlands, Scotland, Sweden, Switzerland | |
| 6 | 18.0 | 10 | Belgium, Corsica, England, France, Germany, Hungary, Italy, Netherlands, Russia, Sweden | |
| 3 | 28.5 | 2 | England, Spain | PPP1CB |
| 6 | 8.0 | 2 | Finland, Scotland | GHITM |

| Chromosome | Position (MB) | Number of outliers | Outlier comparisons | Candidate genes |
|---|---|---|---|---|
| 1A | 70.5 | 2 | Czech Republic, Russia | CDKN1B |
| 4A | 12.0 | 2 | England, Scotland | COL4A5 |
| 3 | 28.0 | 1 | Spain | MRPL33 |
| 6 | 7.5 | 1 | Scotland | BMP1RA |
| 8 | 7.5 | 1 | Netherlands (Vlieland) | |
| 8 | 8.0 | 1 | Belgium | |
| 1A | 48.5 | 1 | Finland | BMP1RA |

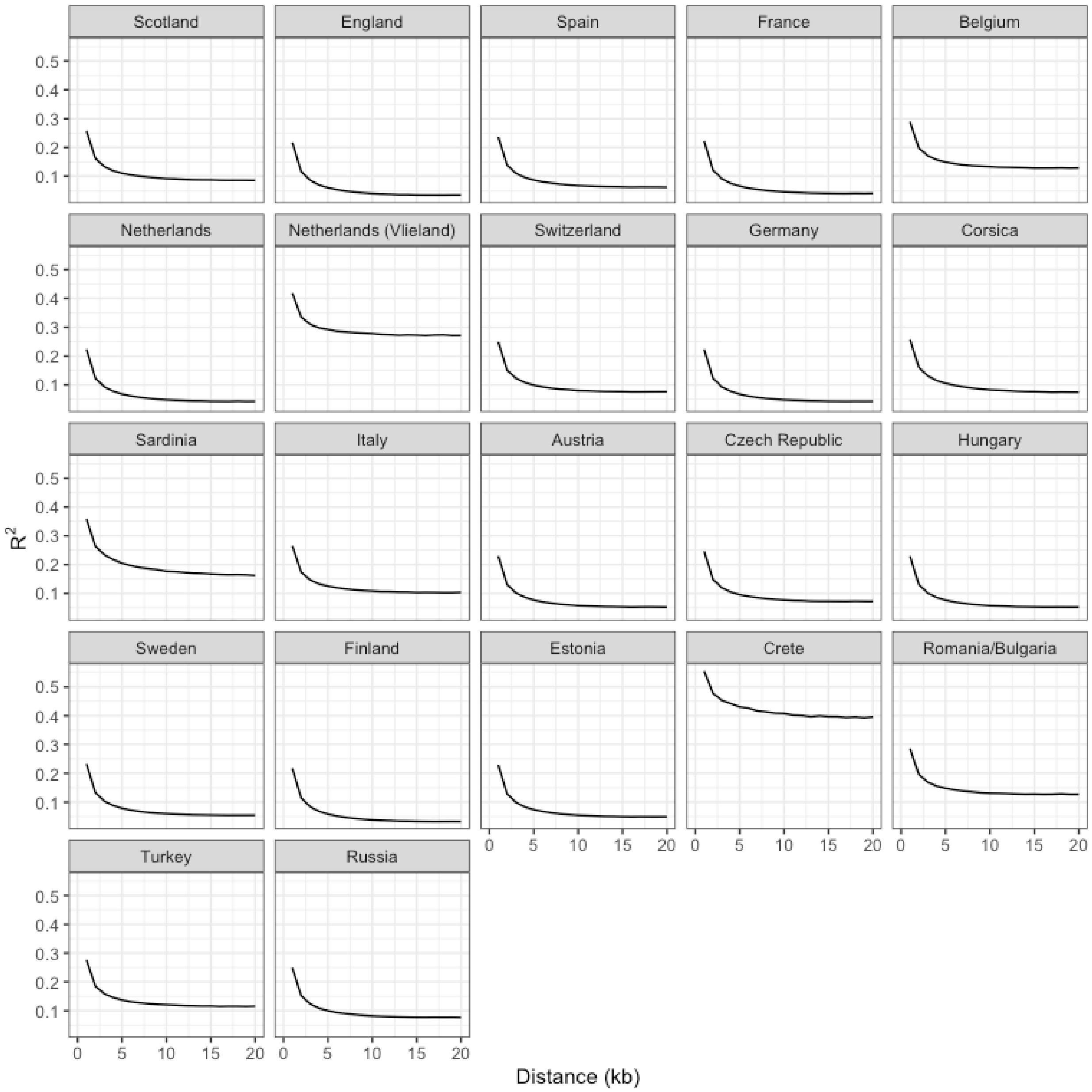


**Figure S1** Linkage disequilibrium in European great tit populations. Lines are means of $R^2$ values for all pairs of markers within 1000bp distance bins.

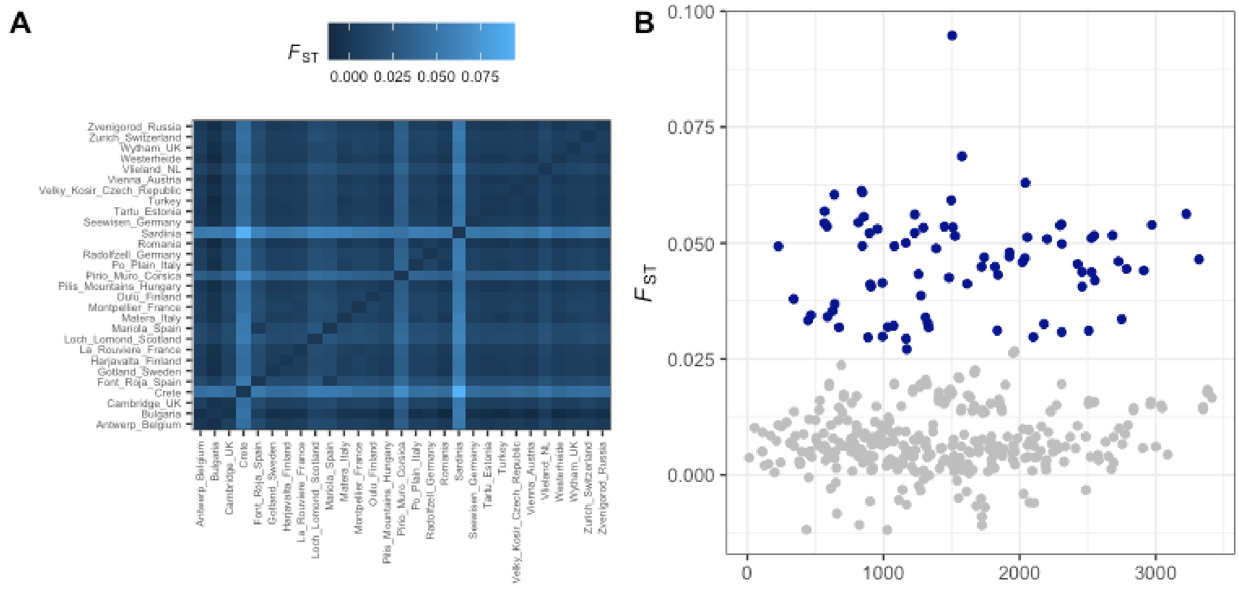


**Figure S2 A** Heatmap of pairwise $F_{ST}$ values between all populations, and **B** Pairwise $F_{ST}$ in relation to geographic distance among European great tit populations. In **B**, pairwise comparisons involving the island populations of Corsica, Sardinia and Crete are coloured in dark blue.

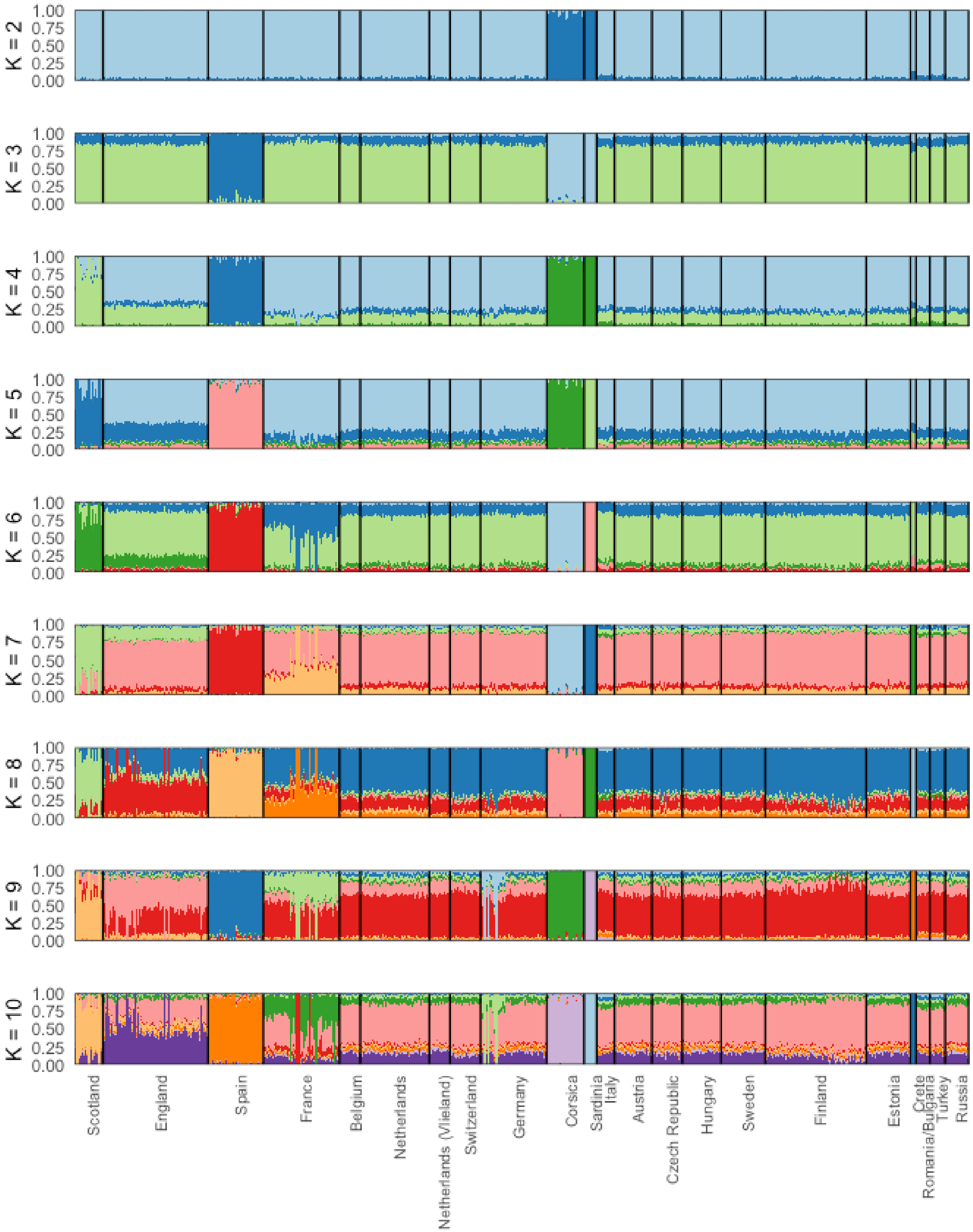


**Figure S3** Admixture analyses at $K$ = 2 to 10 in European great tit populations.

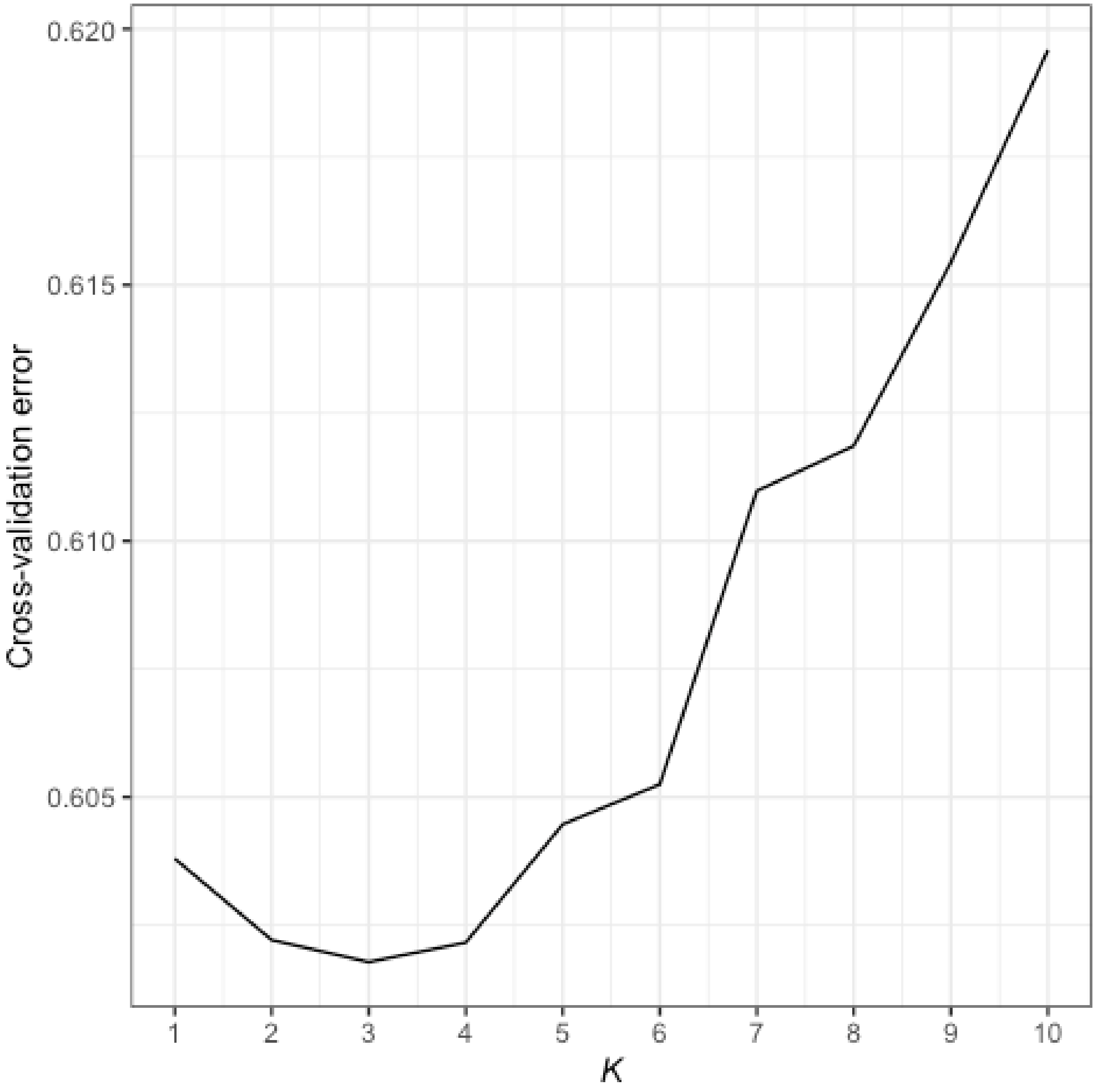


**Figure S4** Cross-validation error from Admixture analyses at $K$ = 1 to 10.

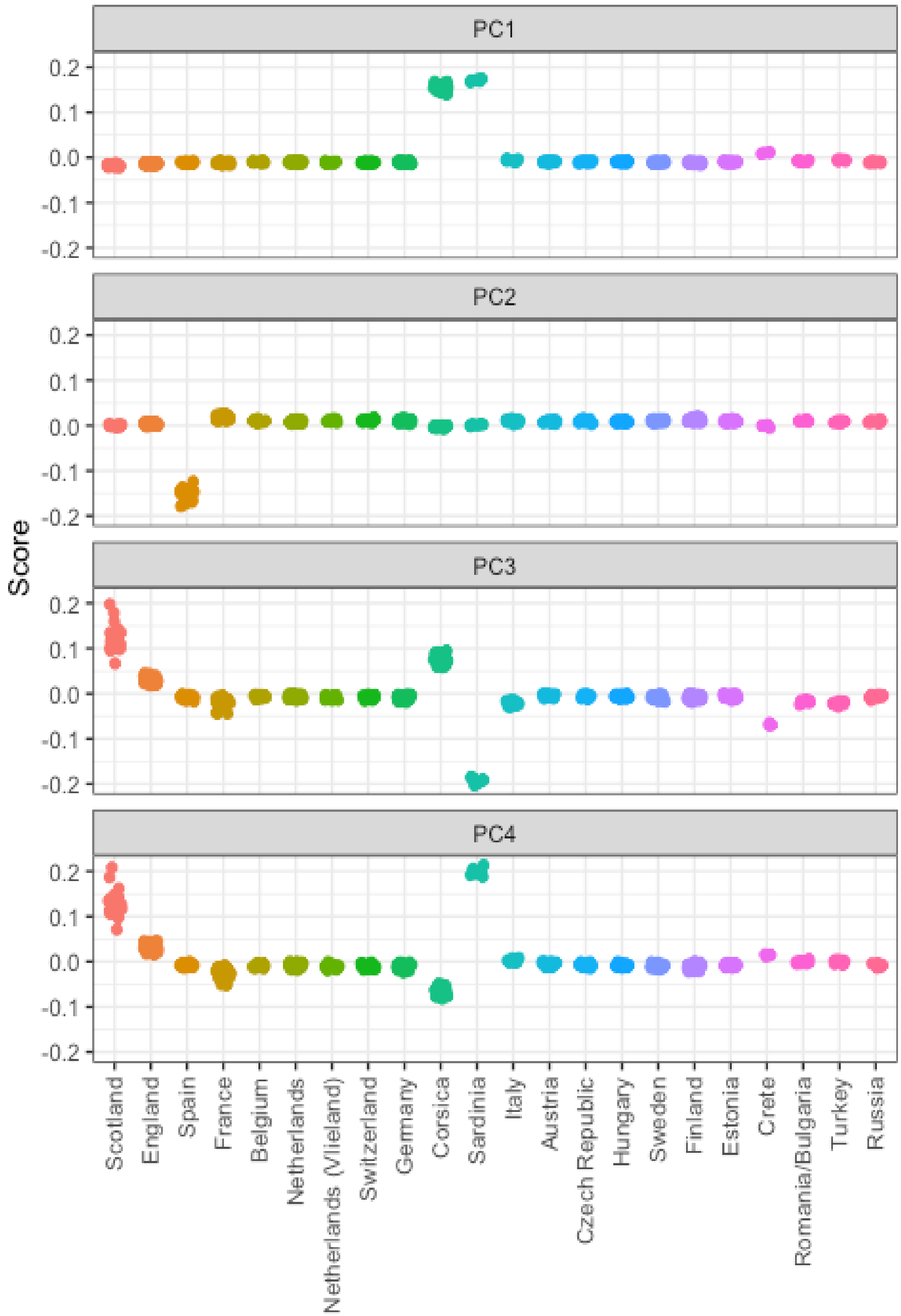


**Figure S5** Results from PCA analysis of European great tit populations.

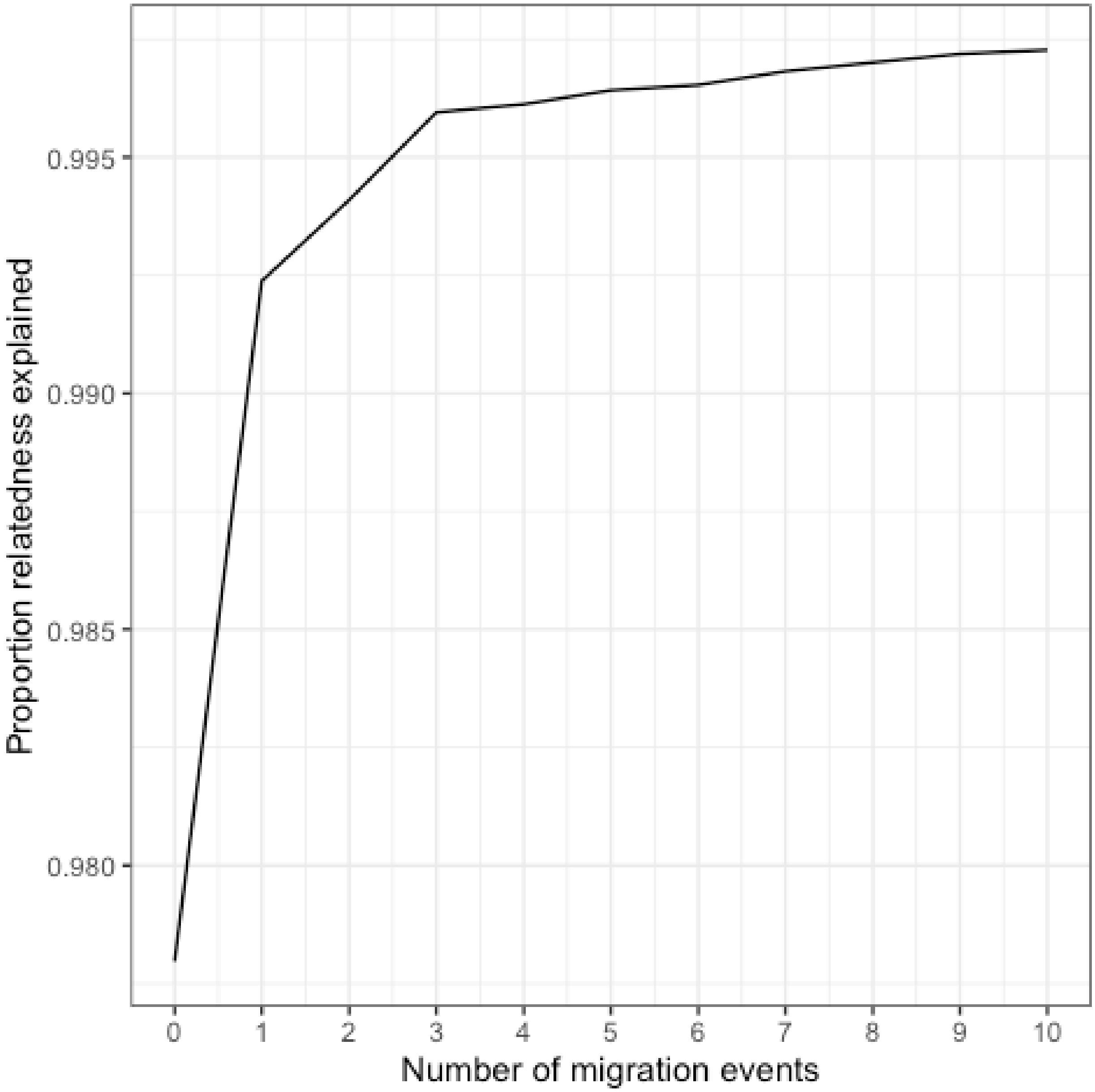


**Figure S6** Proportion of relatedness explained among populations in relation to the number of migration events allowed in TreeMix analyses.

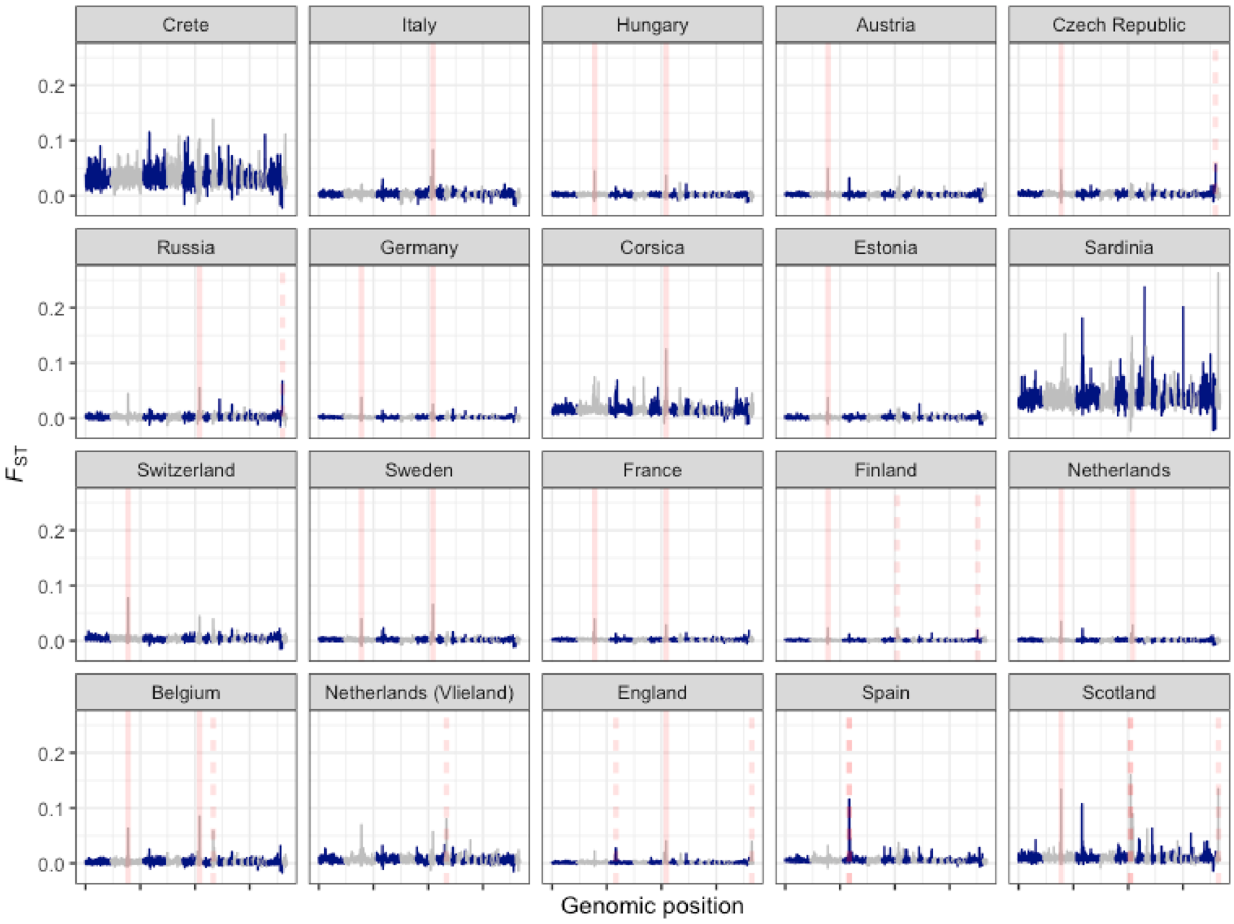


**Figure S7** Landscapes of genomic differentiation in European great tit populations. $F_{ST}$ across the genome is averaged in 500kb windows, with each panel displaying a pairwise comparison with the proposed refugial population in Turkey. Red lines represent $F_{ST}$ outliers (windows with mean $zF_{ST}$ > 10) shared across more than two comparisons (solid red lines), or unique to one or two comparisons (dashed red lines).

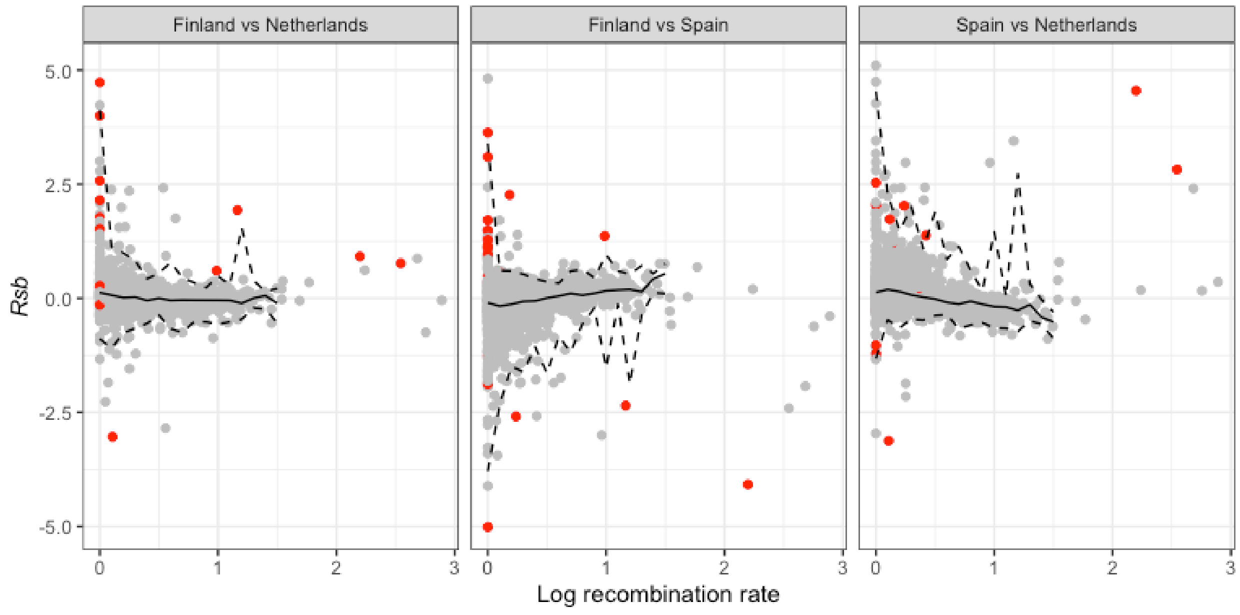


**Figure S8** *Rsb* in relation to local recombination rate in three pairwise comparisons, averaged across 500kb windows. Red points represent the top 1% of FST outliers, and solid and dotted lines represent median and 99% quantiles of *Rsb* windows from bins of 0.1 log cM/Mbp. *Rsb* outliers greater than zero indicate selection in the first population and values less than zero indicate selection in the second population.

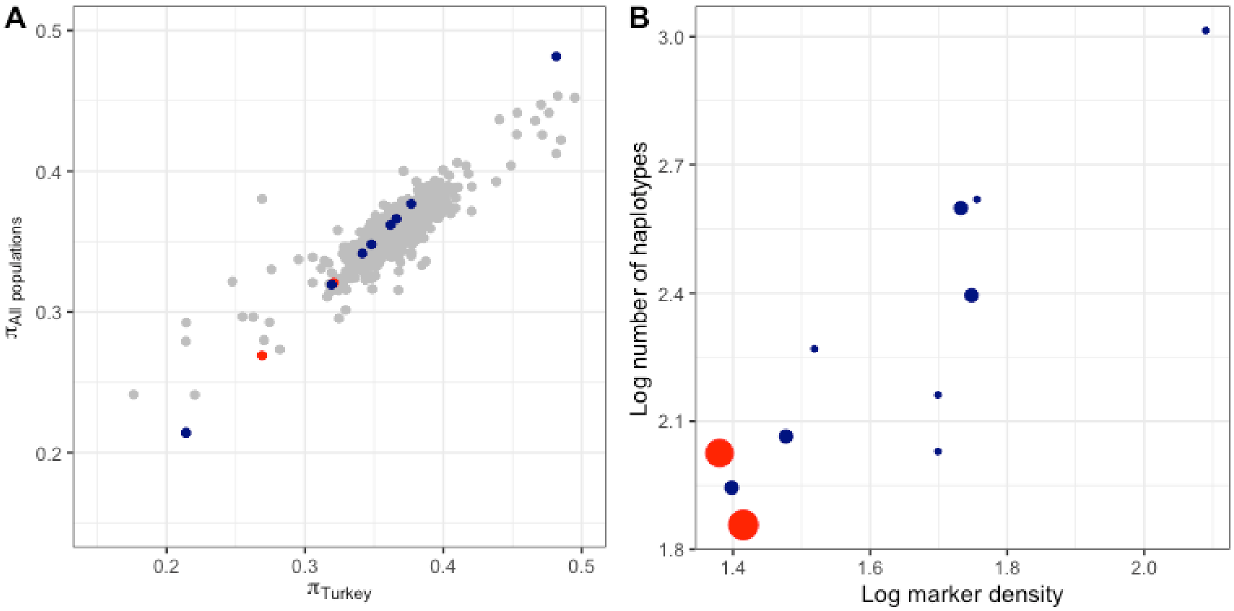


**Figure S9** Genetic diversity in $F_{ST}$ outlier regions in European great tit populations. Each point is an outlier window, with point size scaled by the number of populations the outlier was found in - "shared" haplotypes are coloured red and "unique" haplotypes dark blue (see main text for details). In **A**, Nucleotide diversity for each window is calculated for the proposed refugial population in Turkey, and plotted against the mean for all other populations. In **B**, SNP marker density is plotted against haplotype richness for each region. Low marker density regions tend to have low local recombination rates. Thus, shared regions are more likely to be regions of low recombination rate but unique outlier regions are not.

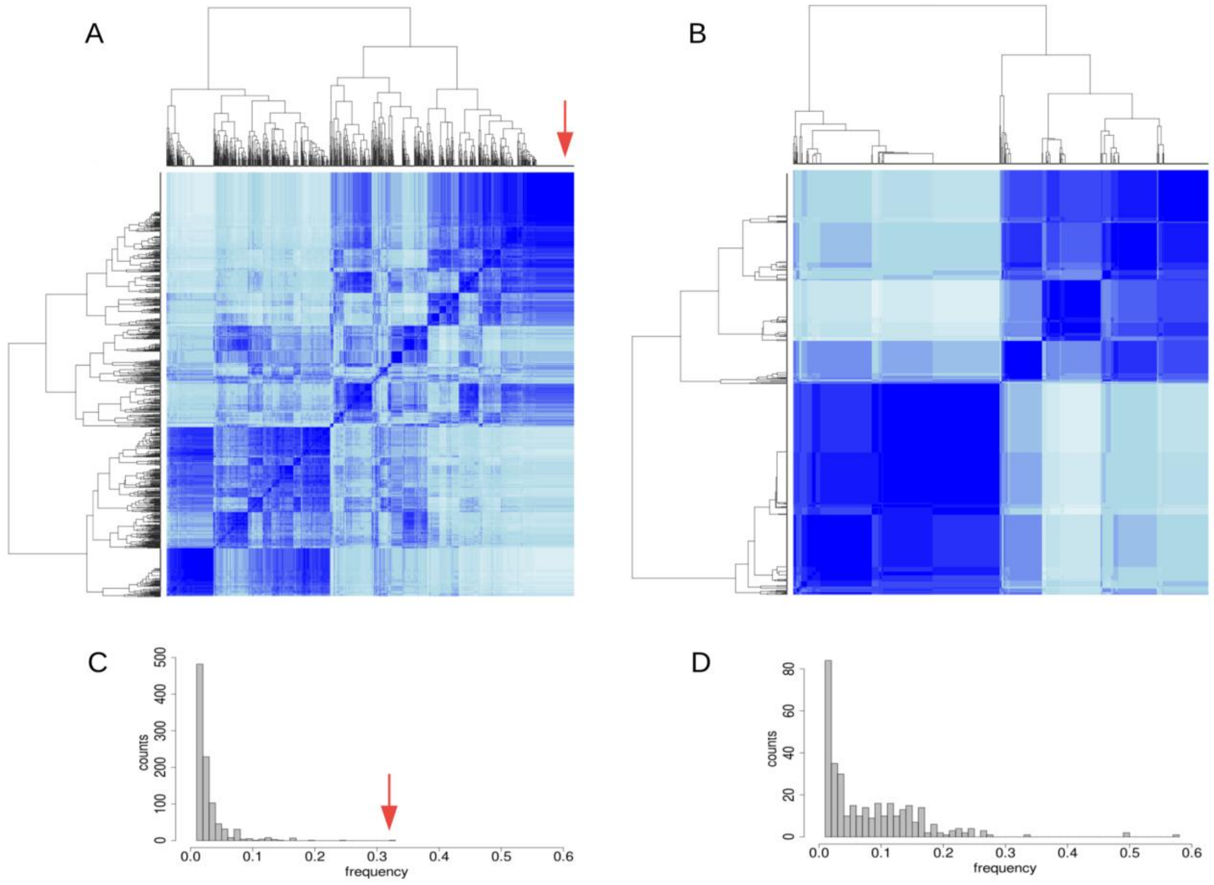


**Figure S10** Haplotype structure underlying regions of high differentiation in a unique vs shared outlier region. **A** IBS clustering of all European haplotypes in the region containing a signal of high differentiation with Turkey specific to the Oulu population. The red arrow indicates the haplotype at highest frequency in Finland. Colour indicates IBS similarity (range 0: white - 1: dark blue). **B** IBS clustering of all European haplotypes in the region on chromosome 2 containing a signal of high differentiation with Turkey in multiple populations. **C** Frequency of all identified haplotypes within each population. The red arrow indicates the high-frequency haplotype within Finland. **D** Frequency of all identified haplotypes spanning the region on chromosome 2 within each population.
